# Impaired brain glymphatic flow in a rodent model of chronic liver disease and minimal hepatic encephalopathy

**DOI:** 10.1101/173526

**Authors:** Anna Hadjihambi, Ian F. Harrison, Natalia Arias, Rocío Gallego-Durán, Patrick S. Hosford, Nathan Davies, Abeba Habtesion, Mark F. Lythgoe, Alexander V. Gourine, Rajiv Jalan

**Author notes:** Joint first authors: A.H and I.H are joint first authors. Joint senior authors: A.V.G. and R.J. are joint last authors. **Contact information:** Rajiv Jalan, Professor of Hepatology, Liver Failure Group ILDH, Division of Medicine, UCL Medical School, Royal Free Campus, Rowland Hill Street, London, NW3 2PF, UK.; <u>or</u>, Alexander V. Gourine, Centre for Cardiovascular and Metabolic Neuroscience, Neuroscience, Physiology and Pharmacology, University College London, Gower Street, London, WC1E 6BT, UK. **Conflict of interest statement:**, **Anna Hadjihambi:** N/A, **Ian F. Harrison:** N/A, **Natalia Arias:** N/A, **Rocío Gallego-Durán:** N/A, **Patrick Hosford:** N/A, **Nathan Davies:** N/A, **Abeba Habtesion:** N/A, **Mark F. Lythgoe:** N/A, **Alexander V. Gourine:** N/A, **Rajiv Jalan:** Rajiv Jalan has research collaborations with Ocera and Takeda and consults with Ocera and has received speaking fees from Norgine. Rajiv Jalan is the inventor of OCR-002, which has been patented by UCL and licensed to Ocera Therapeutics. He is also the founder of Yaqrit limited, a spin out company from University College London. **Author contributions to manuscript: Anna Hadjihambi and Ian F. Harrison:** Performed and analysed the MRI and ICP experiments. Wrote the paper and constructed the figures. **Natalia Arias:** Performed and analysed behavioural experiments. **Rocío Gallego-Durán:** Performed statistical comparisons and figures for behavioural experiments. **Patrick Hosford:** Critical interpretation of data. **Nathan Davies:** Performed bile duct ligation surgery. **Abeba Habtesion:** Performed bile duct ligation surgery. **Mark F. Lythgoe:** Contributed funding to the study, developed the MRI methods, and supervised research staff. **Alexander V. Gourine and Rajiv Jalan:** Led and funded the study. Supervised the manuscript and figure composition. **Abbreviations** HE: Hepatic encephalopathy CSF: Cerebrospinal fluid ISF: Interstitial fluid BDL: Bile duct ligation CLD: Chronic liver disease mHE: Minimal HE Gd-DTPA: Gadolinium ICP: Intracranial pressure ROI: Region of interest SEM: Standard error mean.

## Abstract

Neuronal function is exquisitely sensitive to alterations in extracellular environment. In patients with hepatic encephalopathy (HE), accumulation of metabolic waste products and noxious substances in the interstitial fluid of the brain may contribute to neuronal dysfunction and cognitive impairment. In a rat model of chronic liver disease, we used an emerging dynamic contrast-enhanced MRI technique to assess the efficacy of the glymphatic system, which facilitates clearance of solutes from the brain. We identified discrete brain regions (olfactory bulb, prefrontal cortex and hippocampus) of altered glymphatic flow, which aligned with cognitive/behavioural deficits. Although the underlying pathophysiological mechanisms remain unclear, this study provides the first experimental evidence of impaired glymphatic clearance in HE.

---

The mechanisms underlying the pathogenesis of hepatic encephalopathy (HE) in patients with cirrhosis are not completely understood. Literature suggests that in HE, substances such as lactate, neurosteroids, and bile acids accumulate in the cerebral interstitial and cerebrospinal fluid (ISF/CSF)^1^. The prevailing hypothesis implicates metabolic and signalling deficits induced by hyperammonemia, inflammation and alterations in blood brain barrier function^2, 3^

The lymphatic system is responsible for ISF clearance, a critical mechanism, which maintains tissue homeostasis. Cerebral metabolic rate is very high, accounting for ~20% of body energy expenditure. Although neuronal function is sensitive to alterations in the extracellular environment, until recently, the brain was believed to be devoid of a lymphatic-like clearance system^4^. Recently identified is a brain-wide paravascular pathway that facilitates clearance of various molecules, including toxic interstitial proteins (e.g. (β-amyloid), lactate and others^4^. Subarachnoid CSF enters brain parenchyma along paravascular spaces surrounding penetrating arteries, exchanging with the ISF, facilitating clearance of solutes via convective bulk flow. ISF eventually reaches CNS lymphatic vessels^5^ and systemic circulation. This pathway has been termed the “glymphatic system” due to its dependence on glial mechanisms. Failure of this system may have important consequences and has been linked to the pathogenesis of neurodegenerative disease(s)^6,4^. Here we tested the hypothesis that glymphatic clearance is impaired in HE, potentially contributing to the development of its neurochemical and neurological phenotype.

Dynamic contrast-enhanced MRI was used to visualize brain-wide subarachnoid CSF-ISF exchange in anesthetized bile duct ligated rats (4-weeks; BDL); a model of chronic liver disease (CLD) with minimal HE (mHE)^7^, and sham-operated control animals. Liver dysfunction was confirmed by biochemical and clinical analysis (STable 1). Parenchymal distribution of the MRI contrast agent, gadolinium (Gd-DTPA) throughout the brain was assessed, as it reports the efficacy of the glymphatic flow^1^. Following intra-cisternal infusion of gadolinium, serial acquisition of T1-weighted MR images was performed (Figure 1A).

**Figure 1:**
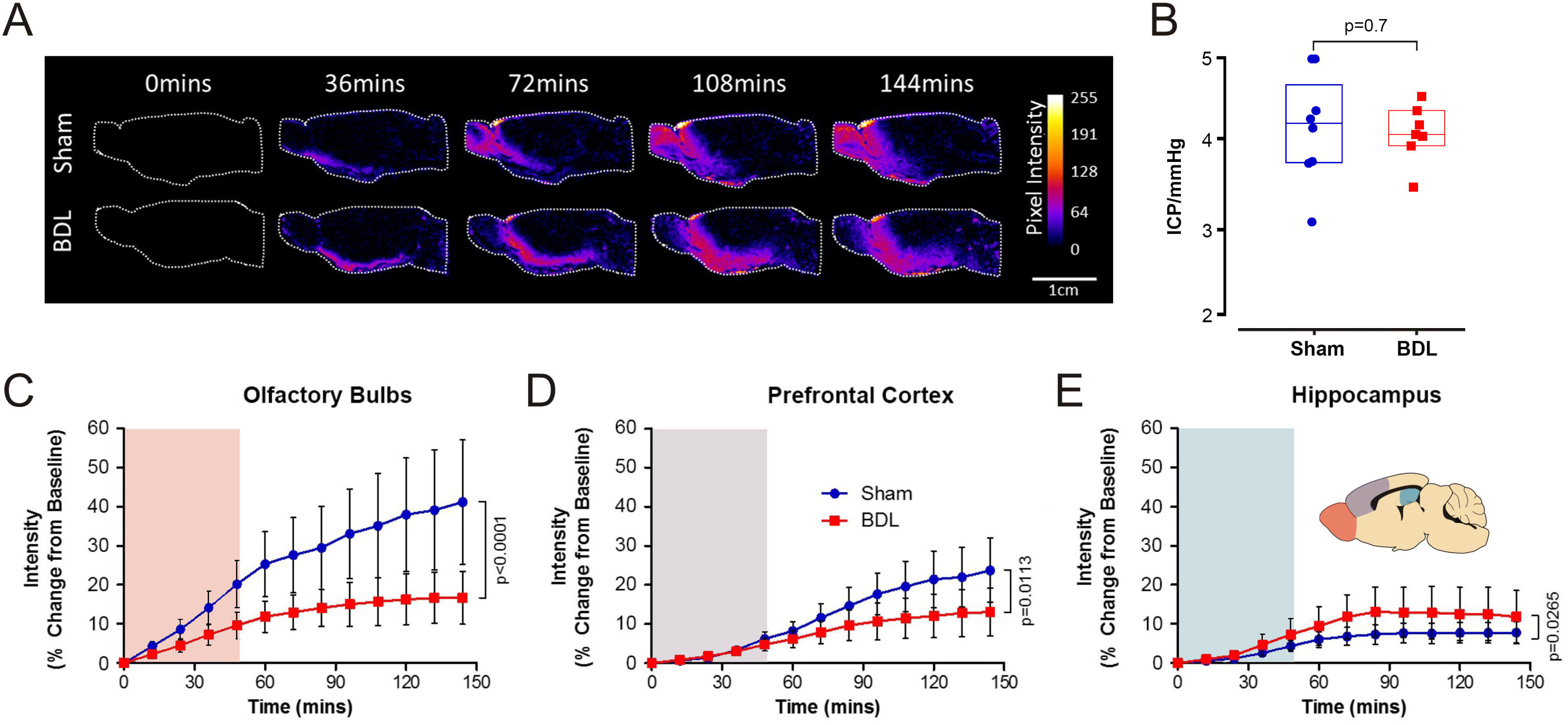
Impaired brain glymphatic flow in HE. **(A)** Representative images of dynamic contrast-enhanced MRI of sham-operated and bile duct ligated (BDL) animals. Pseudocolour scaling illustrates distribution of gadolinium throughout the brain over the 144 min of the recording, with the BDL brain showing reduced glymphatic inflow in rostral areas. **(B)** Summary data illustrating resting intracranial pressure (ICP) in sham-operated and BDL animals. Summary data showing MR contrast intensity changes in **(C)** the olfactory bulb **(D)** the prefrontal cortex and, **(E)** the hippocampus of sham-operated and BDL rats. ***Inset:*** Schematic drawing illustrating the brain regions of interest. Shading indicates period of contrast agent infusion. *p* values indicate the level of significant differences between the sham-operated and BDL groups.

In order to assess the ability of the brain to distribute the infused contrast agent, the brain was compartmentalized (on analysis) and quantification of signal intensity *vs.* time of inflow was compared between the experimental and control groups. CSF filled compartments, including the aqueduct, third and lateral ventricles, showed no differences in flow between sham-operated and BDL animals (SFigure 2A-D). Time-course of the parenchymal distribution of the agent in the striatum, midbrain, caudal cortex, thalamus and hypothalamus was not different between the groups (SFigure 1A-E).

Parenchymal penetration of the contrast agent in the olfactory bulb (p<0.001) and prefrontal cortex (p=0.01) was significantly reduced in BDL compared to sham-operated animals, indicating compromised efficacy of glymphatic flow in these areas (Figure 1C-D). Increased accumulation of contrast agent was observed in the hippocampus of BDL rats (p=0.03) (Figure 1E), further suggesting that altered glymphatic flow in this model of mHE. There were no significant differences in intracranial pressure (Figure 1B), volumes of different brain regions (SFigure 2D), or arrival time of contrast agent (data not shown) between the groups, indicating comparable contrast agent delivery. Therefore, CSF distribution of MRI contrast agent was unlikely to account for the observed differences.

To determine the possible functional associations of altered glymphatic flow, we performed an array of behavioural and cognitive tests in the Barnes Maze. Prefrontal cortex is critically involved in cognitive functions such as working memory, a process of maintaining an active representation of information^8^. We assessed the function of prefrontal cortex by testing spatial working memory. There was no difference between the animal groups in time required to reach the escape box during the sample (training) trial (p=0.3). However, significant differences between the groups in the retention trial were observed (p=0.02) (Figure 2A-B), suggesting that the BDL animals are unable to retain an active representation of information indicative of prefrontal cortex dysfunction.

**Figure 2:**
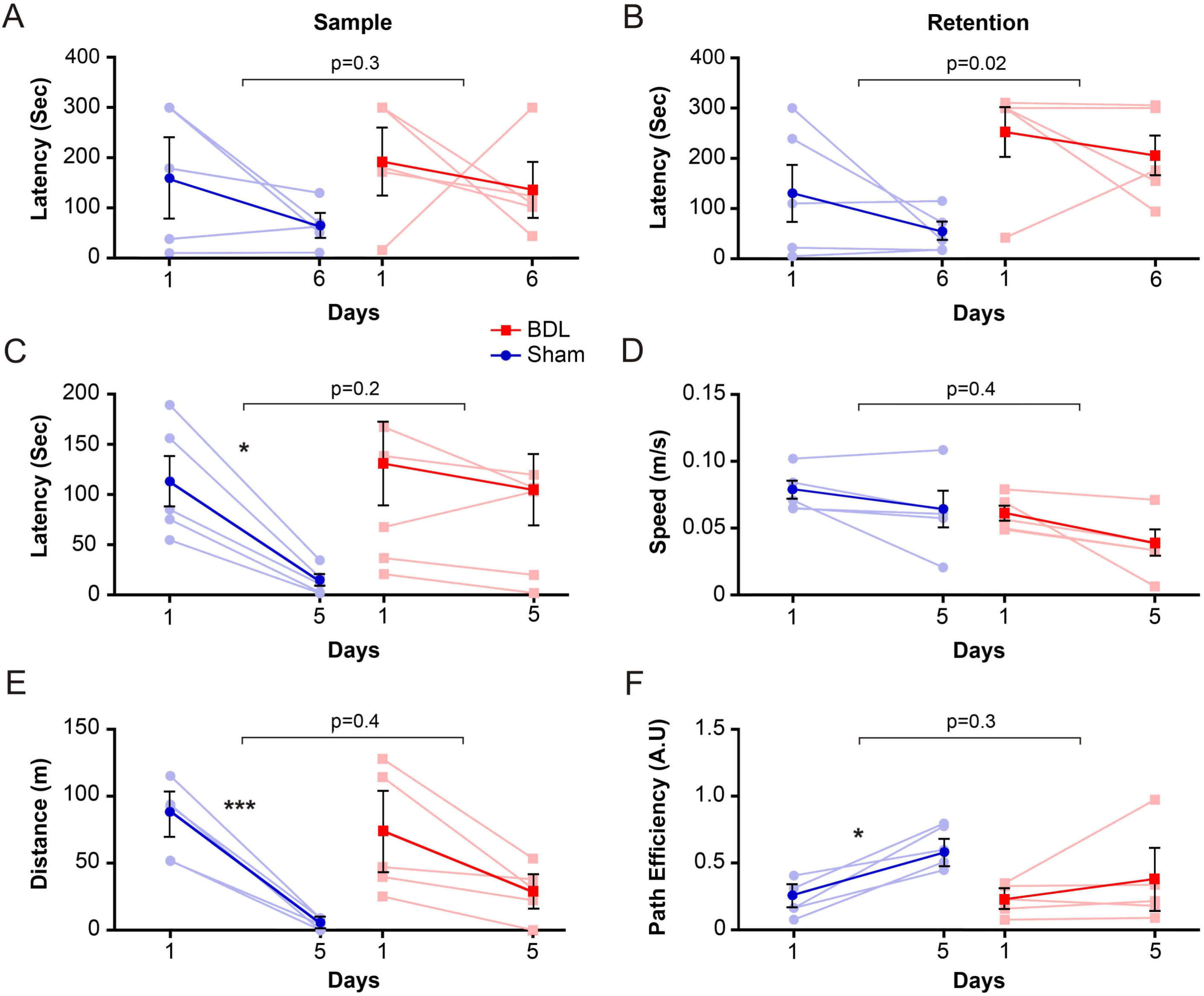
Cognitive/behavioural deficits in HE. (**A**) Summary data illustrating time latency required to reach the escape box in the working memory task in sham-operated and bile duct ligated (BDL) rats. Summary data illustrating **(B)** retention latency in the working memory task, **(C)** latency, in the spatial reference memory task, (**D**) speed during the reference memory task, (**E**) distance travelled during the reference memory task and (**D**) path efficiency from the starting point to the escape box during the reference memory task comparing sham and BDL rats. ***p*** values indicate the level of significant differences between the sham-operated and BDL groups. For within group comparison ^⋆^p<0.05, ^⋆⋆⋆^p<0.001.

As the hippocampus is important in acquisition of spatial reference memory^9^, further tests to assess its function were performed. Although no differences were observed in escape latencies between the groups (p=0.2), the controls showed progressive and significant improvement in the task acquisition during the training days (p=0.04) (Figure 2C). This indicates the inability of BDL animals to learn how to use the reference cues in order to reach the escape box, suggesting impairment of hippocampal function. There were no differences in the speed and distance travelled by animals from the two groups, therefore differences in latency were not due to locomotor deficiency (Figure 2D-E). Moreover, the learning ability of sham-operated, but not BDL animals, was evident from significant decreases in distance covered, which was also reflected in the increase of path efficiency (Figure 2F). These results demonstrate a functional correlate to the observed alteration in glymphatic flow in the hippocampus.

This study demonstrates impaired CSF penetration and parenchymal clearance of the contrast agent in the olfactory bulb and prefrontal cortex, indicating a dysfunction of the glymphatic clearance system in an animal model of mHE. The reasons underlying increased contrast agent inflow in the hippocampus are currently unclear, but may be due to cell loss/neurodegeneration, also seen in models of Alzheimer’s disease or regional glial alterations that may occur in HE^10^. Cytotoxic edema and energy depletion, known features of HE, enhance glymphatic CSF influx while suppressing ISF efflux^11^ may also contribute to the increased contrast inflow. The data showing memory deficits in this model of HE provide a functional correlate for these findings indicating impairment of the prefrontal cortex function as well as a potential derangement in hippocampal-prefrontal cortex connections. The pathophysiological mechanisms underlying impaired glymphatic flow reported here are unclear. However, several factors, some known to be deranged in HE, such as reactive astrogliosis^6^, hemichannel dysfunction^3^, loss of perivascular astroglial aquaporins polarization^12^, arterial pulse-pressure^13^, inflammation^14^, may play a role. In conclusion, this study provides the first experimental evidence of impaired glymphatic clearance in HE.

## Acknowledgements

This study was supported by Grand Challenges UCL and The Wellcome Trust (to Alexander V. Gourine). Alexander V. Gourine is a Wellcome Trust Senior Research Fellow (Refs: 095064 and 200893). Mark F. Lythgoe receives funding from the EPSRC (EP/N034864/1); the King’s College London and UCL Comprehensive Cancer Imaging Centre CR-UK & EPSRC, in association with the MRC and DoH (England); UK Regenerative Medicine Platform Safety Hub (MRC: MR/K026739/1); Eli Lilly and Company. Authors would like to thank Ozama Ismail for his help with the surgical preparation associated with the dynamic contrast-enhanced MRI, Jack Wells for his assistance in setting up the MRI protocol and Christina Elia for her help in producing high quality images.

## Figure legends

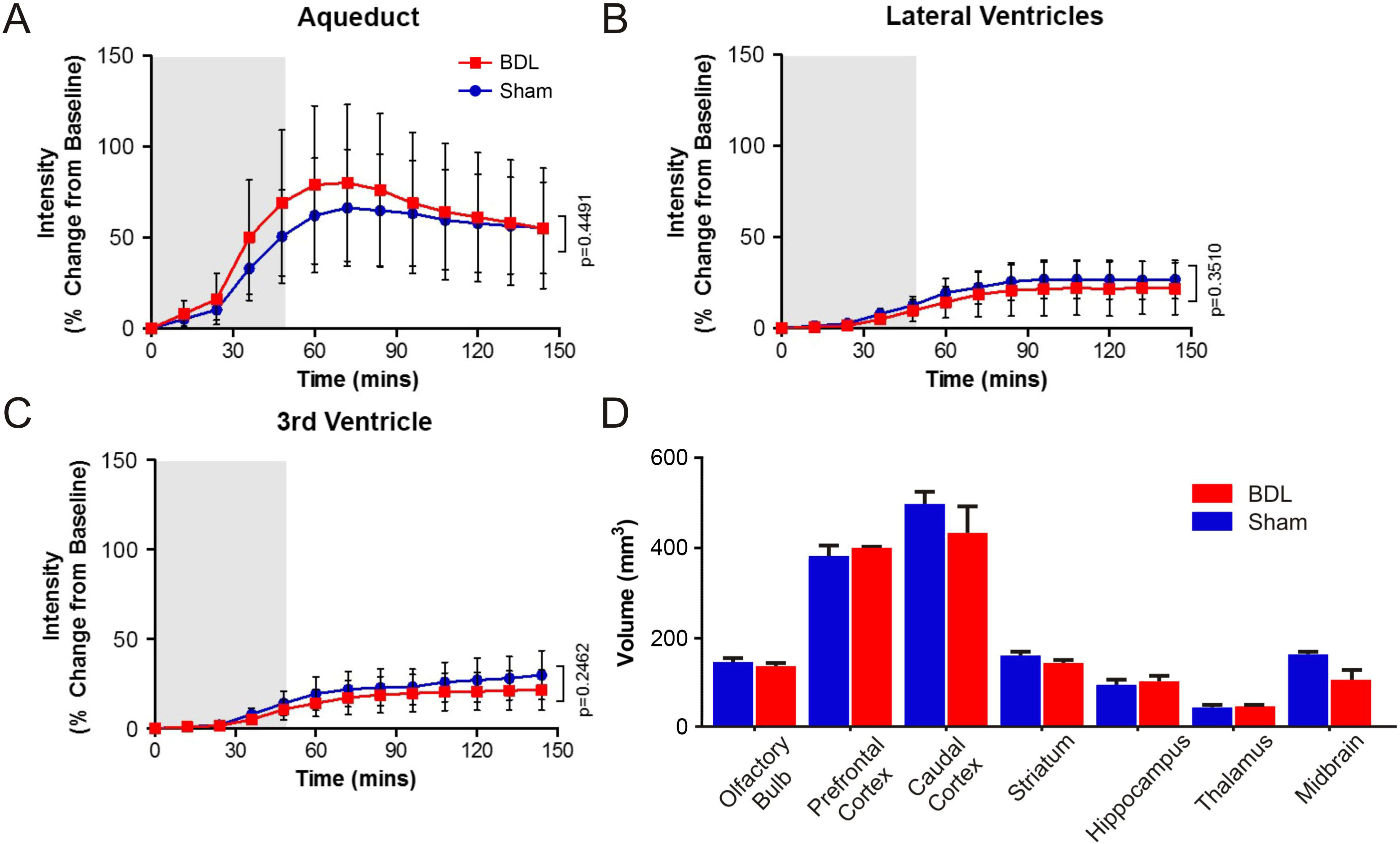

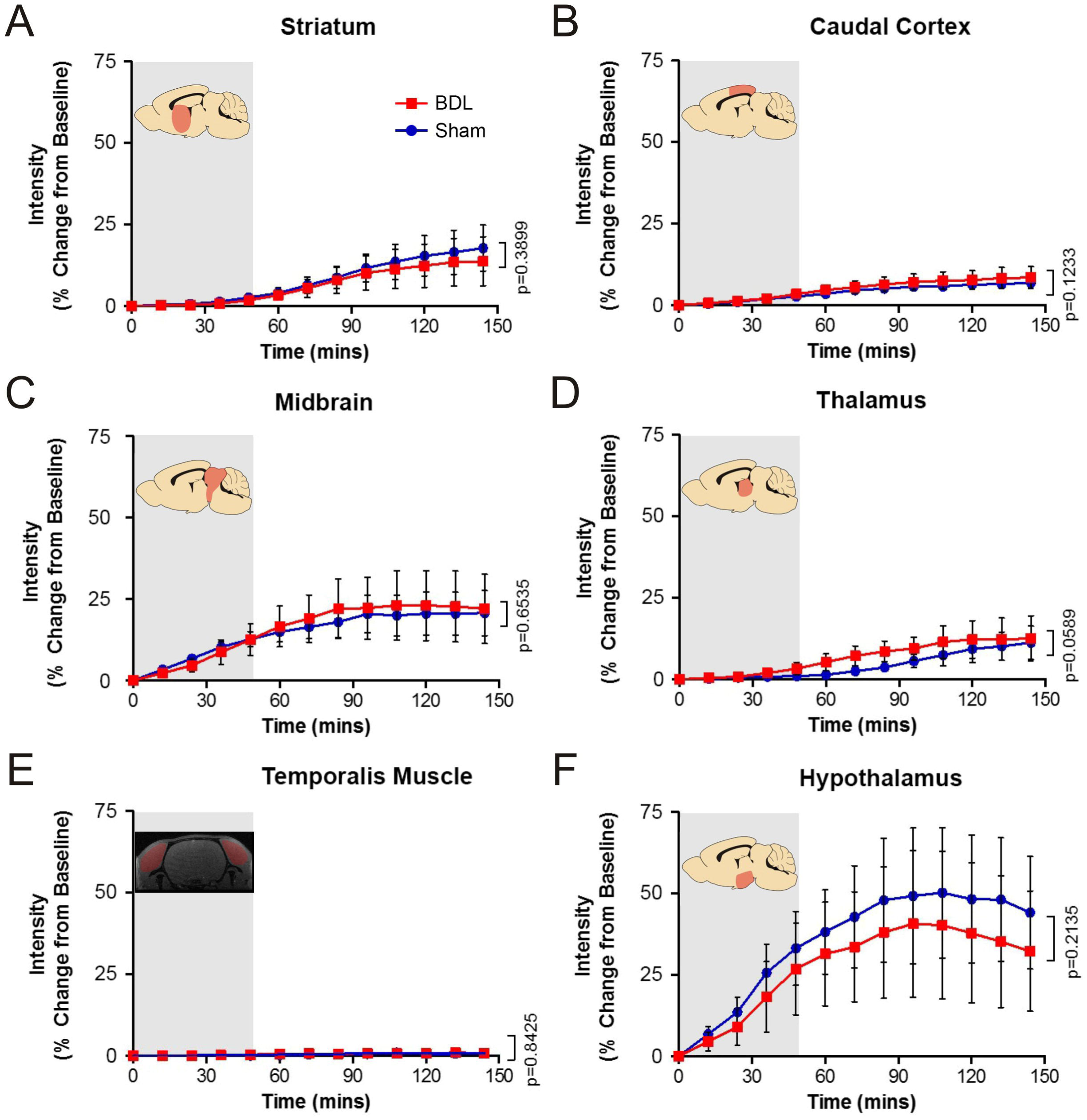

